# Regression models for partially localized fMRI connectivity analyses

**DOI:** 10.1101/2023.04.20.537694

**Authors:** Bonnie B. Smith, Yi Zhao, Martin A. Lindquist, Brian Caffo

## Abstract

Brain functional connectivity analysis of resting-state functional magnetic resonance imaging (fMRI) data is typically performed in a standardized template space assuming consistency of connections across subjects. This can come in the form of one-edge-at-a-time analyses or dimension reduction/decomposition methods. Common to these approaches is the assumption of complete localization (or spatial alignment) of brain regions across subjects. Alternative approaches completely eschew localization assumptions by treating connections as statistically exchangeable (for example, using the density of connectivity between nodes). Yet other approaches, such as hyperalignment, attempt to align subjects on function as well as structure, thereby achieving a different sort of template-based localization. In this paper, we propose the use of simple regression models to characterize connectivity. To that end, we build regression models on subject-level Fisher transformed regional connection matrices using geographic distance, homotopic distance, network labels, and region indicators as covariates to explain variation in connections. While we perform our analysis in template-space in this paper, we envision the method being useful in multi-atlas registration settings, where subject data remains in its own geometry and templates are warped instead. A byproduct of this style of analysis is the ability to characterize the fraction of variation in subject-level connections explained by each type of covariate. Using Human Connectome Project data, we found that network labels and regional characteristics contribute far more than geographic or homotopic relationships (considered non-parametrically). In addition, visual regions had the highest explanatory power (i.e., largest regression coefficients). We also considered subject repeatability and found that the degree of repeatability seen in fully localized models is largely recovered using our proposed subject-level regression models. Further, even fully exchangeable models retain a sizeable amount of repeatability information, despite discarding all localization information. These results suggest the tantalizing possibility that fMRI connectivity analysis can be performed in subject-space, using less aggressive registration, such as simple affine transformations, multi-atlas subject-space registration, or perhaps even no registration whatsoever.

## 1 Introduction

Functional connectivity is defined as the undirected association between two or more brain regions (Friston, 2011). This is often assessed by computing the correlation coefficient over time between spatially different regions of the brain. Most functional connectivity studies employ what we here refer to as a *complete localization approach*. That is, they assume a consistent alignment of regions across subjects. This approach assumes consistent spatial alignment (geography) implying functional alignment. Moreover, it is pursued using data transformed into a common template space (Talairach and Tournoux, 1988; Mazziotta et al., 1995).

A shortcoming of this approach is that a key assumption underlying it is often violated, namely that brain regions are functionally aligned across subjects. However, the presence of large inter-subject differences in functional localization is known to remain even after structural alignment to a standard template (Haxby et al., 2014). This problem can be partially addressed using spatial smoothing at a cost in spatial resolution, effect size, and power. Alternatively, it can be addressed using analytic approaches such as Hyperalignment (HA, Haxby et al., 2011, 2014, 2020). HA is a functional alignment technique that attempts to register individual brains based on functional properties rather than only on anatomical locations. It seeks to harness variation in functional connectivity to create a common functional space. Brain data from local regions are iteratively mapped into a common high-dimensional space using a Procrustes transformation, which preserves and aligns participants based on local representational geometry. This procedure has been shown to increase functional similarities across subjects while preserving subject-specific information (Haxby et al., 2011; Guntupalli et al., 2016; Feilong et al., 2018; Nastase et al., 2019).

In contrast, in recent work, Tang et al. (2023) relaxed the complete localization assumption to its limit, by treating intra-subject connectivity between regions as if exchangeable. This assumption of exchangeability (see Draper et al., 1993, for an example) implies the use of the empirical distribution of connections as a sufficient statistic, thus completely ignoring spatial information. This “connectivity distribution” has interesting properties, including being theoretically robust to registration. In addition, this approach is useful in settings, such as in stroke, surgical, or degenerative disease settings, where localization assumptions are both suspicious and difficult to employ because of registration challenges (Scheinost et al., 2012) (see Tward et al., 2020, for a discussion of registration in non-standard settings). However, such exchangeability assumptions lie at an extreme, ignoring the well-established core neuro-organizational principle of common localized functional specialization. In comparison, functional HA approaches and related functional alignment techniques (Xu et al., 2012; Andreella et al., 2022; Wang et al., 2022) fall somewhere in between full spatial localization and connectivity exchangeability.

In this work, we propose a new approach that also lies between these two extremes. However, the proposed approach differs a great deal from functional alignment since: (i) explicit alignment is not a goal of the analysis; (ii) aggregate connectivity effects are considered under the assumption that they are exchangeable within levels of covariates; and (iii) the approach can be implemented entirely in subject space. For this latter point, we envision a use for this technique when using multi-atlas registration, where a single, common, template is not employed and instead template information is carried from a collection of labeled atlases to the subject (Rezende et al., 2019).

Instead of alignment as a goal, we use region-pair (or voxel-pair) information to explain within-subject connection strength using linear models, as is typically done in network analyses (Salter-Townshend and McCormick, 2017). Our models treat subject-level Fisher-transformed regional connection matrices as the outcome of interest and use geographic distance, homotopic distance, network labels, and region indicators as covariates to explain variation in connections. This style of analysis allows us to summarize the composition of the fraction of variability explained by each of these characteristics of the region pairs. Additionally, we investigate data repeatability (Bridgeford et al., 2021; Airan et al., 2016; Wang et al., 2021b; Finn et al., 2015) under our proposed approach. Also known as test-retest reliability, data repeatability quantifies the consistency over time of multiple measurements made on the same subject. Bridgeford et al. (2021) proposed the data repeatability metric of *discriminability*, which is the probability that two measurements for the same subject will be more similar than measurements for two different subjects. We compare discriminability for resting-state fMRI data from the Human Connectome Project (HCP, Van Essen et al., 2013) under our approach, and under the completely localized and the non-localized approaches.

The paper is organized as follows: in Section 2, we describe the data, present our models, and describe how we assess variability explained and discriminability. In Section 3, we present results for the HCP data, and Section 4 concludes with a discussion.

## 2 Methods

### 2.1 Data from the Human Connectome Project

The dataset consists of resting-state fMRI data from 470 healthy subjects from the Human Connectome Project (HCP, Van Essen et al., 2013) 500 subject release. All data were acquired on a Siemens Skyra 3T scanner at Washington University in St. Louis. Subjects completed two fMRI sessions on consecutive days. Each session included two 15-minute resting-state scans, one with a right-to-left and the other with a left-to-right phase encoding. In this work, we focus solely on the left-to-right phase encoding data, and hence our data consists of 1200 brain volumes (TR=720ms) for each day. Further description of the data and processing pipelines applied can be found in (Geuter et al., 2018). Briefly, scans were preprocessed using the HCP “fMRIVolume” pipeline (Glasser et al., 2013), which includes gradient unwarping, motion correction, fieldmap-based distortion correction, brain-boundary-based registration to structural T1-weighted scan, non-linear registration into MNI152 space, grand-mean intensity normalization, and spatial smoothing using a Gaussian kernel with a full-width half-maximum of 4 mm. This was followed by time series extraction using the Shen atlas of 268 regions of interest (Shen et al., 2013) via regional means. The Fisher’s Z transform was taken of the synchronous temporal correlations across regions, resulting in 268 choose 2, or 35,778 transformed inter-regional correlations for each subject and session.

### 2.2 Subject-level connectivity regression models

We begin by defining notation. Let us denote the Fisher’s Z transformed empirical correlations over time between two regions *j* and *j*′ by *Z*(*j, j*′), for *j, j*′= 1, …, *R*, where *R* is the number of regions or seed locations (*R* = 268 in our case). Let *Z* be the symmetric matrix comprised of the *Z*(*j, j*′), with zeros on the diagonal, and let *Ƶ* = {*Z*(*j, j*′)}_*j<j*′_ be the collection of *R* choose 2 correlations viewed as a set. We note that each of these refers to a subject- and session-specific measurement. When considering multiple subjects and sessions, we will write *Z*_*i,t*_(*j, j*), *Z*_*i,t*_, and *Ƶ*_*i,t*_ to denote measurements for subject *i*, session *t*.

In a localized approach to functional connectivity, subject-specific correlations are compared across a common alignment of regions. For example, edge-wise regression approaches regress *Z*_*i*_(*j, j*′) on subject-specific covariates, for each edge (*j, j*′). By contrast, a completely non-localized analysis treats the elements of *Ƶ* as exchangeable, and thus characterized by their order statistics or empirical distribution function.

As an alternative to these approaches, we consider a subject-level connectivity regression model on the subject-specific collection of Fisher transformed correlations. Consider a loss function that minimizes ∑_*j<j*′_{*Z*(*j, j*′) − *X*(*j, j*′)′*β*}^2^ over *β*, where *X*(*j, j*′) is an ℝ^*p*^-vector containing an intercept term and region characteristics, and where the parameter *β* is an ℝ^*p*^-vector of regression coefficients. Let *C*_*j*_ = (*c*_1*j*_, *c*_2*j*_, *c*_3*j*_) be the voxel coordinates of the center of mass of region *j*. Further, assume that a value *m* in the first coordinate represents an approximate mid-sagittal plane. We consider the following predictors:

1. A spline basis for homotopic distance between regions *j* and *j*′, if the two regions lie in different hemispheres. Thus, the spline is based on the distance {(*c*_1*j*_ + *c*_1*j*′_ − 2*m*)^2^ + (*c*_2*j*_ − *c*_2*j*′_)^2^ + (*c*_3*j*_ − *c*_3*j*′_)^2^}^1*/*2^*I*{Sign(*m* − *c*_1*j*_) ≠ Sign(*m* − *c*_1*j*′_)}. This allows for the investigation of the impact of approximate symmetry without the need for registration to a symmetric template. The rationale for this is that homotopic correlations are among the most reproducible resting state findings (Zhao et al., 2022a).
2. A spline basis for geographic distance between regions *j* and *j*′, ||*C*_*j*_ − *C*_*j*′_||. Geographic correlations can arise from biologically irrelevant reasons, such as processing (smoothing), as well as biological reasons, such as functional specialization.
3. Intra- and inter-network membership indicators. Network information was based on the results of Finn et al. (2015), who group the 268 regions in the Shen parcellation into eight networks. For each network *k*, we include an indicator for whether both regions *j* and *j*′ are in network *k* (8 intra-network terms), and for each pair *k < k*′, we include an indicator that one of regions *j, j*′ is in network *k* and the other is in network *k*′(28 inter-network terms).
4. Indicators of regional involvement. For each region *r*, we include an indicator that *r* is region *j* or region *j*′, *I*(*j* = *r*) + *I*(*j*′= *r*) − *I*(*j* = *r*)*I*(*j*′= *r*). (Thus, *R* = 268 such terms were included.) The rationale for these terms is the possibility that specific regions may exhibit consistently higher or lower connectivity.

For the terms involving spline bases, we used generalized additive models (GAMs Hastie, 2017; Wood, 2011) with thin plate regression splines. The inclusion of region and network terms introduced rank deficiency, so that (as usual) terms were dropped until a full rank design matrix was obtained.

Here, we have fit these models on data that has been transformed to a common template space (see Section 2.1). However, we emphasize that this style of analysis could also be conducted in subject-space (under the modification of not including network terms). For example, it could be used in multi-atlas settings where labeled templates are registered to subjects, rather than subjects being morphed into a common template space. In particular, homotopic distance and geographic distance only require subject-level information, such as identifying an approximate mid-sagittal plane. Therefore, we refer to our connectivity regression approach as being “partially localized”.

It is also worth emphasizing that this approach is a form of matrix regression (Zhao et al., 2021, 2022b). Specifically, the regression estimates for *β* are the same as those obtained by minimizing the norm 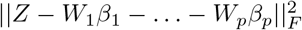, where the *W*_*k*_ are matrices with element (*j, j*′) equal to element *k* of *X*(*j, j*′) and *β*_*k*_ is element *k* of *β* and || · ||_*F*_ is the Frobenius norm (that is, the square root of the sum of the squares of the matrix elements).

To summarize, the characteristics used included: non-linear terms for geographical distance, non-linear terms of homotopic (symmetric inter-hemispheric) distance, separable region effects, and network effects.

### 2.3 Partitioning proportion of variability explained

A substantial benefit of the proposed model is that it allows us to investigate the relative importance of each of the characteristics described above. For linear models, Grömping (2006) has presented a number of relative importance metrics, which are implemented in the `relaimpo` R package. The importance of a given predictor can be assessed using the amount by which the coefficient of determination *R*^2^ increases when that predictor is added to the model. However, this amount will typically depend on whether other predictors have already been included in the model. To account for this, the recommended `lmg` metric of Grömping (2006) averages the increase in *R*^2^ over all possible orderings of the predictors. In this way, a given predictor’s relative importance weighs both the direct impact of that predictor on *R*^2^ when it is the only predictor in the model and the indirect impact when it is added after one or more of the other predictors. Here we adapt this for our setting, where we focus on the relative importance of groups of predictors in a generalized additive model (with Gaussian family and linear link). Specifically, we consider all network terms as one group, all separable region effects as a second group, and geography and homotopy as the third and fourth groups of predictors, respectively. We then use the increase in *R*^2^ when a given group of predictors is added, and average over possible orderings of the four groups. For each subject-specific model, this partitions the total proportion of variation explained by the model into the proportion of variation explained by each of these four groups of predictors. This provides a means of comparing relative importance of different types of predictors within a given model. (Note though that we cannot necessarily compare values across different models.) When comparing the relative importance of these groups of predictors, we should note the unequal sizes of the groups, with the group of region effects containing by far the largest number of terms, followed by the group of network terms.

We additionally considered a measure of average connectivity for each region and each pair of networks based on our connectivity regression model. Because the regression coefficients for region effects and for network effects are to some extent aliased for each other, we do not focus on the magnitude of each individual coefficient. Instead, for each region *j*, we consider the linear predictor from our model for each pair of regions (*j, j*′), and take the average of these linear predictors as the average connectivity for region *j*. Similarly, we take the average connectivity for the pair of networks (*k, k*′) to be the average of the linear predictors for each pair of regions where one region is in network *k* and the other is in network *k*′. We then average each of these over subjects and sessions.

### 2.4 Repeatability

In addition to an analysis of the features (coefficients) obtained across subjects using our connection-regression framework, the degree of repeatability obtained by summarizing subjects in this manner was also considered via discriminability (Bridgeford et al., 2021). Population discriminability is the probability that two measurements for the same subject will be more similar than two measurements from different subjects, under a given distance metric. To elaborate, let 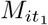 and 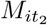 be measurements for the same subject at two different sessions and similarly *M*_*i*′*t*′_ for a different subject at a potentially different session. Let *d*(·, ·) be any distance metric compatible with the measurements. Thus, discriminability (with respect to this distance *d*(·, ·)) is defined as: 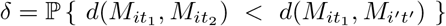, which is well-defined assuming independent subjects and that this probability does not depend on the specific subjects and sessions being considered (Wang et al., 2020). Given *n* independent subjects each measured at *T* sessions, a consistent and unbiased estimator is given by the sample discriminability 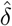 (Bridgeford et al., 2021; Wang et al., 2020):

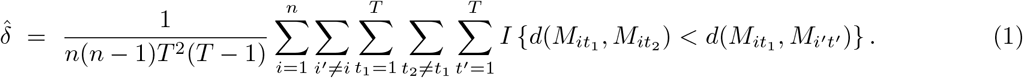

To compute discriminability under a given approach, one must first specify an inter-subject/inter-session distance based on the type of measurements used. In the fully localized approach, given two correlation matrices, *Z*_*i,t*_ and *Z*_*i*′,*t*′_, we use the Euclidean distance between the vectorized strictly upper triangular elements, which is equal to 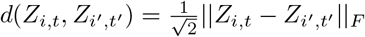.

For our regression approach, we take the measurement to be the matrix 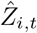 of fitted values (projections) from the regression model, and we use the same Euclidean distance as above, applied to 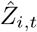. In comparison to the non-localized approach which treats all correlations as exchangeable, our approach treats correlations as exchangeable only within levels of the covariates in our model—a less extreme assumption. We note that this approach to discriminability does use a localized distance to make comparisons across subjects. For the regression models, this distance is exactly a Mahalanobis distance of the coefficients around the variance/covariance matrix of the regressors. A more purely unlocalized approach would simply use the regression coefficients themselves (i.e., a Mahalanobis distance weighted around an identity matrix). Comparing the projections (weighted distance), as opposed to comparing coefficients (unweighted), has the benefit of being fixed in dimension across different models and independent of the distance units used.

In the non-localized approach, each measurement is an unordered set *Ƶ*_*i,t*_, so a Euclidean distance cannot be used with this approach. Instead, we use the 2-Wasserstein distance (see Galichon, 2018) between the connection distributions. This is simply the Euclidean distance between the quantiles of *Ƶ*_*i,t*_ and the quantiles of *Ƶ*_*i*′,*t*′_.

### 2.5 Approaches and Models Compared

Within our connectivity regression approach, we consider four models that use varying degrees of information about the region-pairs as predictors. All models include geographic distance and homotopic distance between pairs of regions. The first model (‘Reference model’) uses only these predictors, and is closest to a non-localized approach. The other three models additionally include network terms (‘Reference model + networks’), separable region terms, (‘Reference model + regions’), or both (‘Reference model + networks + regions’). We investigate the proportion of variation in the HCP data explained by each class of predictor in each of these four models, and we compare discriminability in the HCP data under each of our models and under the localized and non-localized approaches.

## 3 Results

### 3.1 Proportion of variation explained

Figure 1 shows the total variation explained within each subject with the most complex model (‘Reference model + networks + regions’). Across subjects, between 25% and 75% of the variation in correlations is explained by the model. The median proportion of variation explained is over 50%. Figure 2 partitions the subject-level variation explained by each component (color), in each of the partial localization models (panel). Table 1 gives the means over all subjects. Homotopy explained surprisingly little variation and was the lowest with geography second. It is interesting to note that the distribution of explained variation was somewhat insensitive to the model. For example, the distribution for network remains fairly similar whether region is included or excluded. Region was the largest component explaining on average roughly 40% of the variability in the largest model. Note that the number of terms for each type of predictor also plays a role here, since the partition is based on the increase in *R*^2^ when all terms in a given category are added to the model (for example, 268 region terms, or 36 network terms). Here the model ‘Reference model + networks + regions’ was fit after dropping region terms for identifiability. We also re-fit this model by instead dropping network terms for identifiability. As expected, this resulted in slightly more variability explained by region terms (0.41 rather than 0.39) and slightly less variability explained by network terms (0.11 rather than 0.13), with geography and homotopy essentially unchanged.

**Figure 1:**
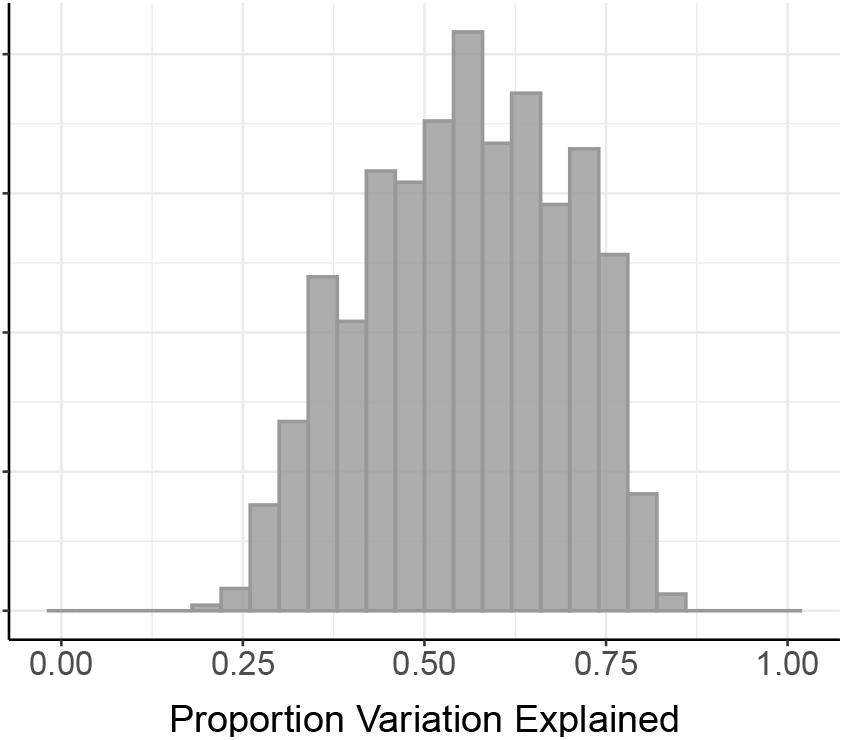
Proportion variation explained. Shown is the total proportion variation explained by a model including geographic, homotopic, network, and region effects, for each of the 470 subjects in the HCP data and each session. The median across all subjects and sessions is 0.56.

**Figure 2:**
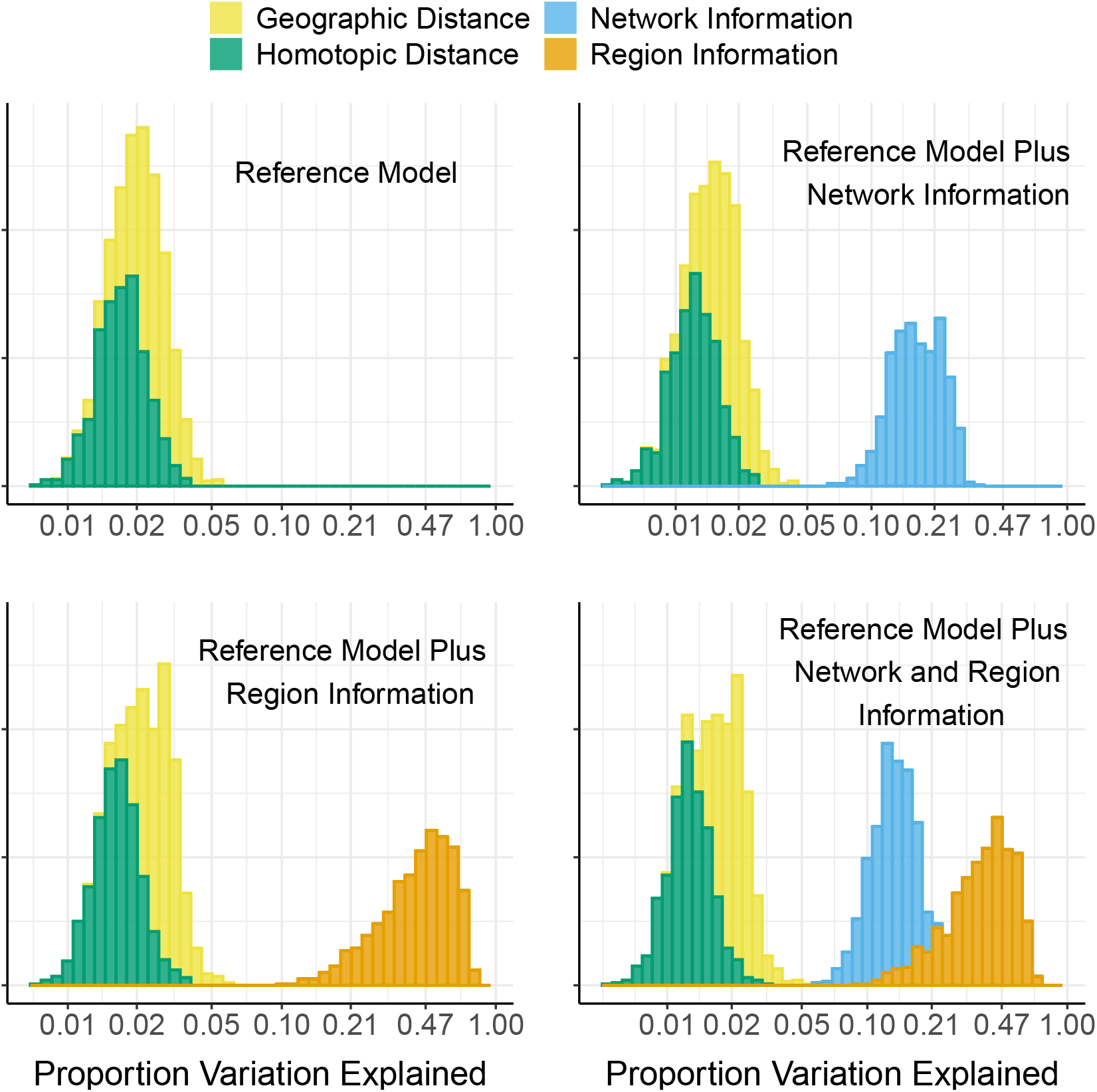
Partition of the proportion of variation explained. Each histogram shows the proportion of variation explained by each type of predictor used in a given model, for each of the 470 subjects in the HCP data. Types of predictors are: geographic distance between pairs of regions, homotopic distance between pairs of regions, (intra- and inter-) network membership indicators, and indicators of involvement for each region. Note that the *x*-axis is linear on the log_10_ scale.

**Table 1:**
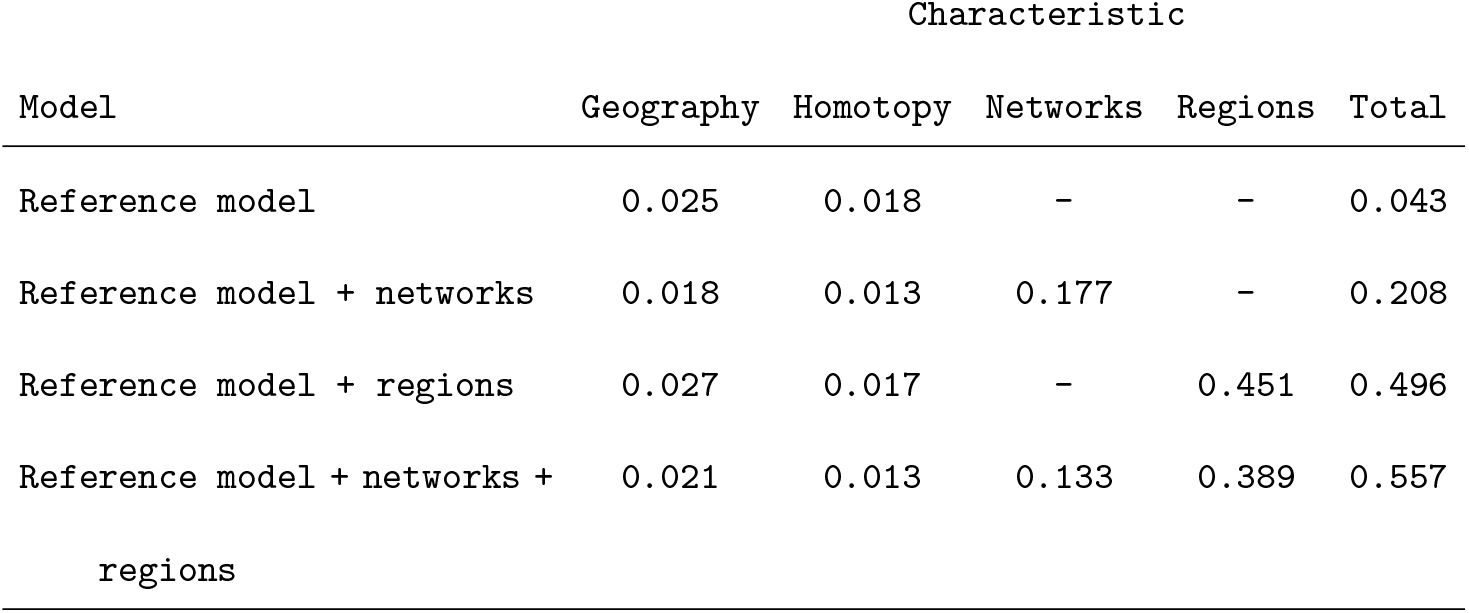
Mean proportion variability explained using the four considered models, averaged over the 470 subjects and two sessions in the HCP data. The reference model contains only an intercept and smooth functional terms for homotopic and geographic distance. The other models also include an indicator for each region, network indicators, or both.

We also assessed, in the ‘Reference model + networks’ model, whether intra-network (*j* and *j*′ belonging to the same network) or inter-network terms (*j* and *j*′ belonging to different networks) account for more variability explained. Figure 3 shows the proportion of variability explained by geography, homotopy, intra-network terms, and inter-network terms; the distributions for intra-network terms and inter-network terms are the same.

**Figure 3:**
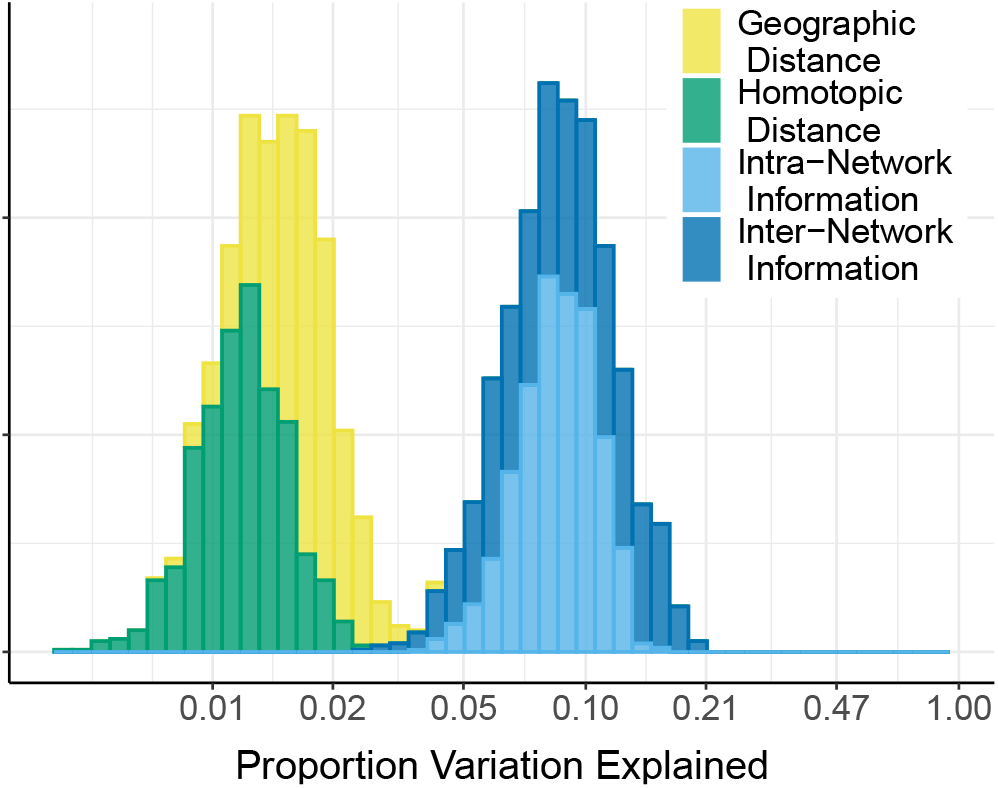
Comparison of intra- versus inter- network effects. Shown is the proportion of variation explained by each type of predictor in a model that included geographic, homotopic, and network effects, for each of the 470 subjects in the HCP data. Types of predictors considered are: geographic distance between pairs of regions, homotopic distance between pairs of regions, intra-network membership indicators, and inter-network indicators. Note that the *x*-axis is linear on the log_10_ scale.

Figure 4 shows those regions with the greatest average predicted connectivity, as described in Section 2.3. Most of these regions are in the motor network or one of the visual networks of Finn et al. (2015). Table 2 shows estimates and 95% confidence intervals for the mean predicted connectivity for each pair of networks. Confidence intervals are bias-corrected and accelerated (BCa) bootstrap intervals (Efron, 1987). As expected, average predicted connectivity tends to be high within a given network, with the subcortical-cerebellum network being the exception. Within-network average connectivity was especially high for the visual networks, and there was also high between-network connectivity among the visual networks, and between the motor network and the visual networks.

**Table 2:**
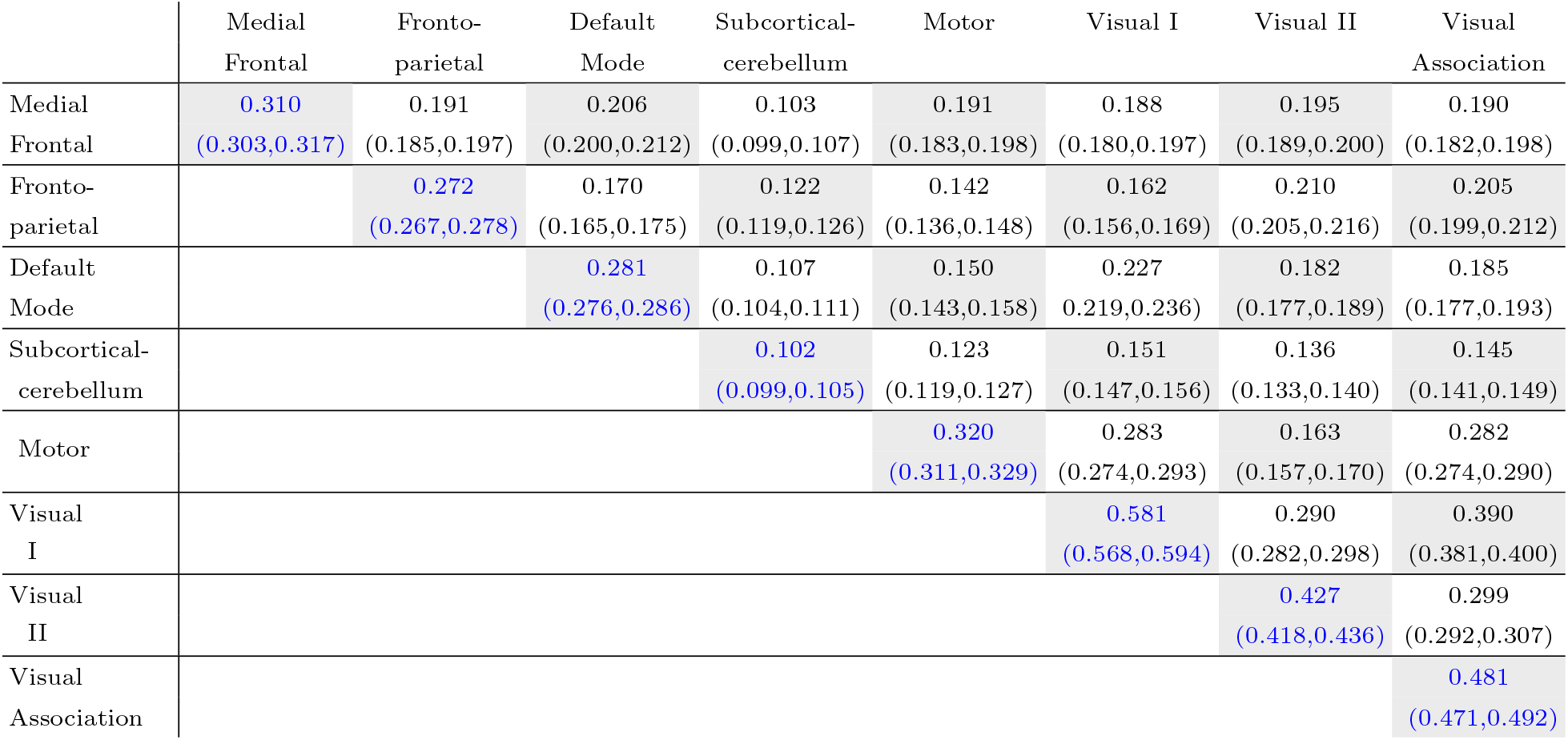
Estimates and 95% confidence intervals of mean predicted correlation for each pair of networks. The estimates in the diagonal entries are the average predicted correlation among pairs of regions which are both in the given network, averaged over subjects and sessions in the HCP data. Estimates in the off-diagonal entries are the average predicted correlation among pairs of regions from the two distinct networks, averaged over subjects and sessions. Confidence intervals are BCa bootstrap intervals from 10,000 resamples of the subject-specific predicted average network connectivities.

**Figure 4:**
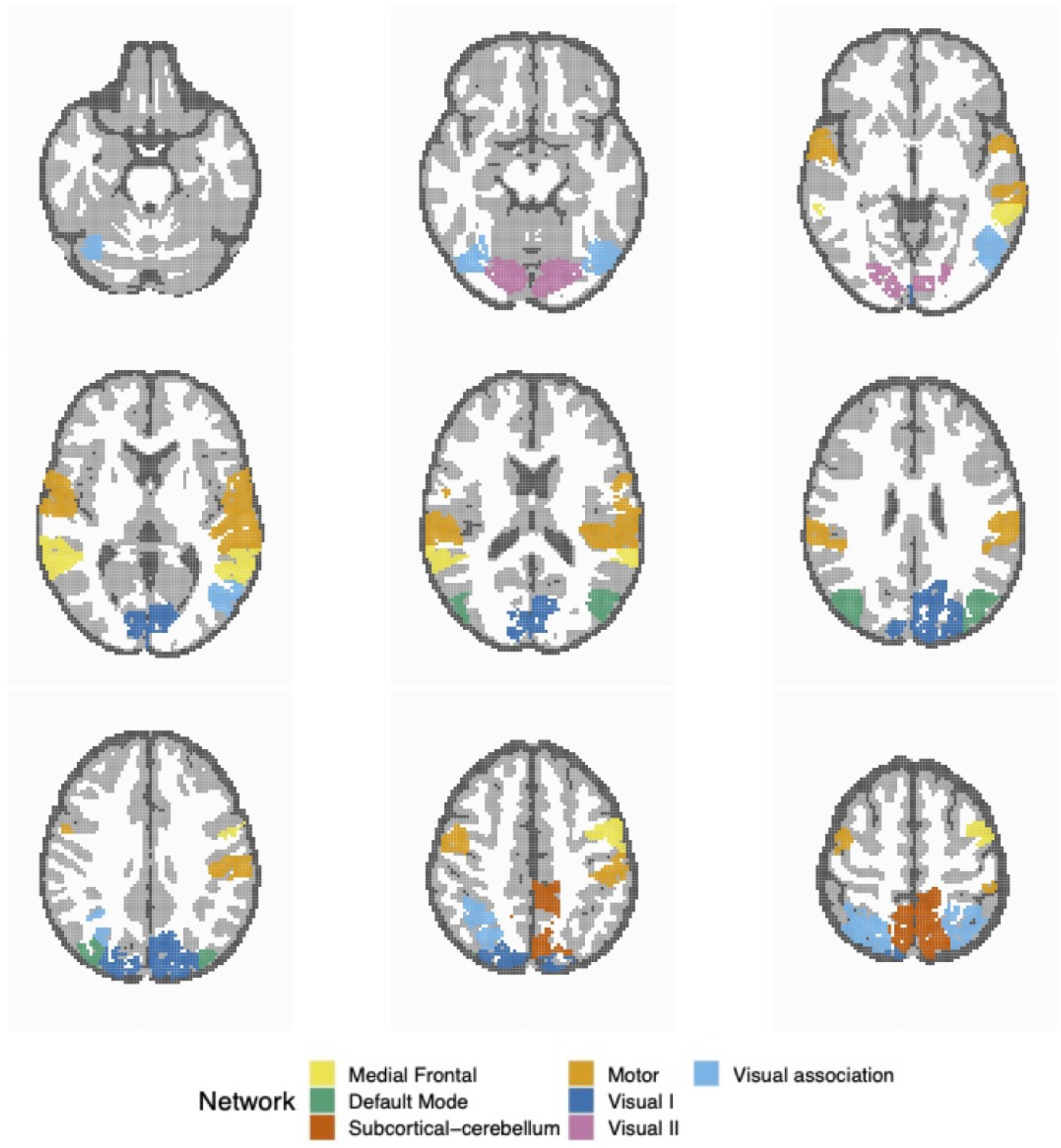
Regions with the largest average predicted correlation, color-coded by network membership. Of the 29 regions shown here, 3 regions are in the Medial Frontal network, 2 are in the Default Mode network, 3 are in the Subcortical-Cerebellum network, 8 are in the Motor network, 5 are in Visual I, 2 are in Visual II, and 6 are in Visual Association. Predictions are from the model including geographic, homotopic, network, and regional terms.

### 3.2 Discriminability

Table 3 shows the point estimate, 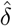, of discriminability under each of the approaches and models described in Section 2.5. Of note, the Wasserstein distance exchangeability approach (completely non-localized) resulted in a fairly high discriminability. In addition, the largest regression model obtains a discriminability that is comparably as high as the fully localized (i.e., full connectivity matrix) approach. It is worth noting, however, that the discriminability compared here puts correlations back in the matrix form, and thus uses a localized distance. We do this primarily for comparability with the fully localized approach. To summarize, subjects are largely identified by even weak summaries of their connectivity matrices.

**Table 3:** Estimates of discriminability for the HCP data under three approaches. The localized approach uses the Euclidean distance of the vectorized Fisher Z transformed connectivity matrices as the distance. The partially localized approach does the same; however, the matrix is the projection of the estimated connections from the linear model. The fully non-localized approach uses the 2-Wasserstein distance on the empirical distribution of the Fisher Z transformed connectivity values.

| Approach |  | Discriminability |
| --- | --- | --- |
| Non-localized |  | 0.657 |
| Partially<br>Localized | Reference Model | 0.658 |
|  | Reference Model + Networks | 0.712 |
|  | Reference Model + Regions | 0.773 |
|  | Reference Model + Networks and Regions | 0.781 |
| Localized |  | 0.860 |

## 4 Discussion

In this paper, we introduce a set of simple subject-level regression models to characterize connectivity, while only partially utilizing localization information. These subject-level models use the Fisher-transformed regional connection matrices as an outcome and geographic distance, homotopic distance, network labels, and region indicators as covariates. We contrast the amount of localization information included in our models with respect to the amount of variation in the connections explained, explanatory power, and data repeatability.

Data repeatability, assessed using discriminability, increased from 66% to 86% depending on whether we used a non-localized, partially localized, or fully localized approach, and on which effects were included in the model. It is interesting that the connectivity distribution approach resulted in a relatively high discriminability, despite the extreme loss of information incurred by considering only the collection of connections. In fact, a model that attempts to explain connectivity using only geographic proximity and homotopy does exactly the same. Adding network membership and node variables gets fairly close to fully localized discriminability. However, these results should be taken with a grain of salt. Data repeatability is a somewhat weak property as it is a statement about measurement variance and distinctness of a trait, regardless of the measure’s utility as a biomarker. For example, images of human fingerprints are highly repeatable but contain no meaningful biological information. In addition, data repeatability is not insensitive to demographics that impact measurements. For example, a dataset with a mixture of older and younger subjects will be more repeatable than one with only subjects from a single age group (Wang et al., 2021b).

With regard to the variability explained, it is not surprising that node-level connectivity and network membership had such large effects. Node-level connectivity has significance related to both biology and nuisance reduction. Biologically, it is hypothesized that specific foci are “hubs” of network activity (see Van Den Heuvel and Sporns, 2011, for example). A benefit of the proposed approach is that it does not require hub locations to be consistent across subjects. From a nuisance perspective, any contaminant that increases or decreases connectivity in a spatially varying manner would receive some benefit from node adjustment in such a regression model. Similarly, the intercept includes both nuisance and possibly real effects, conflating global nuisance effects and increased overall connectivity, and is related to global signal regression (Liu et al., 2017). Network variables are unsurprising as having strong effects since they are somewhat circularly used in this application. That is, networks are exactly sets of nodes that exhibit strong inter-node correlations consistently across subjects (in the sample used by Finn et al., 2015).

The proportion of variability explained by these simple effects ranged from extremely high (75% of the variation in connections) to low (25% of variation in connections). The majority of the variation was explained by nodes and networks, with the largest amount of variation explained by the nodes. Geography explained greater variability than homotopy, both typically less than 5% of the total variation. The low values for homotopy are somewhat surprising, given that bi-laterally symmetric correlations are some of the strongest and most consistent fMRI findings. However, homotopy was very loosely defined as the distance across an approximate mid-sagittal plane. It is possible that the non-linear spline terms do not fully account for homotopic effects. In addition, homotopy (as defined) was aliased with geography for all pairs close to the mid-sagittal plane. Finally, the impact of lateralized networks might have dragged down the impact of homotopy.

A benefit of the proposed style of analysis is that labels could be applied in subject space, allowing for multi-atlas registration. However, performing analyses in template space in this paper allowed us to compare results with a fully localized approach. Future work will investigate how data repeatability is impacted by fitting the connectivity regression models in subject space. It should be mentioned that including network labels does require registration. In addition to networks, we considered models where we leveraged the inter-subject registration to consider common PCA approaches (Wang et al., 2021a). This was done to consider the impact of the specific network estimation tool. Common PCA components had a similarly strong effect to those of the network labels; however we did not use them in our analysis, primarily because of the challenging issue of inter-subject rank selection.

Finally, we note that the analysis approach is not entirely novel. Intra-subject graph estimation combined with inter-subject analyses of graph metrics is a common method of analysis (see Wang et al., 2010), including the evaluation of repeatability (Andellini et al., 2015; Braun et al., 2012). However, weighted graph analyses in a regression model using these terms, along with measuring proportional variation explained, are less explored. We further believe that extending these methods to whole-brain analyses is possible and would lead to novel subject-level summaries of connectivity. Here, the primary issue is computational, since the number of correlations grows at the rate of the number of voxels squared, leading to hundreds of millions of correlations needing to be modeled.

## Acknowledgments

Data were provided by the Human Connectome Project, WU-Minn Consortium (Principal Investigators: David Van Essen and Kamil Ugurbil; 1U54MH091657), which was funded by the McDonnell Center for Systems Neuroscience at Washington University and the 16 NIH Institutes and Centers that support the NIH Blueprint for Neuroscience Research.

